# UniScore, a unified and universal measure for peptide identification by multiple search engines

**DOI:** 10.1101/2024.10.09.617445

**Authors:** Tsuyoshi Tabata, Akiyasu C. Yoshizawa, Kosuke Ogata, Chih-Hsiang Chang, Norie Araki, Naoyuki Sugiyama, Yasushi Ishihama

**Affiliations:** Graduate School of Pharmaceutical Sciences, Kyoto University, Kyoto 606–8501, Japan; Graduate School of Medical Sciences, Kumamoto University, Kumamoto, 860-8556, Japan; Omics Research Center, National Cerebral and Cardiovascular Center, Suita, Osaka 564-8565, Japan; Laboratory of Proteomics for Drug Discovery, National Institute of Biomedical Innovation, Health and Nutrition, Ibaraki, Osaka 567-0085, Japan

**Keywords:** UniScore, Search engine, Peptide identification, Reanalysis, False discovery rate, Chimera spectrum

## Abstract

We propose UniScore as a metric for integrating and standardizing the outputs of multiple search engines in the analysis of data-dependent acquisition (DDA) data from LC/MS/MS-based bottom-up proteomics. UniScore is calculated from the annotation information attached to the product ions alone by matching the amino acid sequences of candidate peptides suggested by the search engine with the product ion spectrum. The acceptance criteria are controlled independently of the score values by using the false discovery rate based on the target-decoy approach. Compared to other rescoring methods that use deep learning-based spectral prediction, larger amounts of data can be processed using minimal computing resources. When applied to large-scale global proteome data and phosphoproteome data, the UniScore approach outperformed each of the conventional single search engines examined (Comet, X! Tandem, Mascot and MaxQuant). Furthermore, UniScore could also be directly applied to peptide matching in chimeric spectra without any additional filters.

## Introduction

The identification of amino acid sequences from peptide tandem mass spectra has been carried out for more than 30 years, and in the early days, *de novo* sequencing was performed by reading the b- and y-ion series that are predominantly observed in low-energy collision-induced dissociation reactions in triple quadrupole and ion trap mass analyzers.(1) In the post-genome era, as protein amino acid sequence databases have become more extensive and very large numbers of spectra have been obtained using LC/MS/MS in the data-dependent acquisition (DDA) mode, a number of search engines have been developed to identify peptides from tandem mass spectra, including Sequest(2) and Mascot(3), which are representative of the search engines that match MS/MS spectra obtained experimentally against databases. In the early days, the threshold for identification was set based on statistical scores specific to each search engine, but after the introduction of the target-decoy (TD) approach(4), identification criteria based on false discovery rate (FDR) became common. As public data repositories as well as various commercial and non-commercial search engines (e.g. X! Tandem(5), OMSSA(6), Andromeda/MaxQuant(7, 8), Byonic(9), MSGF+(10)) were developed, many large-scale studies were conducted and attempts were made to create complete human proteome atlases. The first attempts(11), (12) simply merged the results of a large number of different research groups, so many of the hit proteins were redundant, while the decoy hits piled up, resulting in poor data quality(13). The problems with the target-decoy method were highlighted, and an improvement in the FDR control method was proposed(14). Meanwhile, even without such large-scale data, attempts have also been made to increase the number of hit proteins by integrating the identification results from multiple search engines for a single raw data set. However, if the results of each search are simply merged, the rate of overlap between target hits will be higher than the rate of overlap between decoy hits, and this will result in a higher FDR than that set for each search. As a solution to this problem, a number of tools have been reported that merge the results of different search engines while controlling the FDR(15), (16), (17, 18), (19).

In another method called peptide sequencing tag (PST)(20), proteins are separated and digested, and the sample solution is analyzed by infusion-nanoESI-MS/MS without LC. This requires a prior interpretation step in which the researcher assigns partial amino acid sequences obtained from MS/MS spectra by manual inspection, and in addition to the interpreted sequences, the mass of the precursor peptide, the mass of the first peak in the identified sequence ladder, and the mass of the last peak in the ladder are entered as query data for searching against the protein database. PST is potentially the most powerful search method(21). However, it has rarely been applied to LC/MS/MS data because it is difficult to fully automate the step of prior interpretation of MS/MS spectra.

Search engines such as Sequest, Mascot, X! Tandem and MaxQuant have been applied not only to high-resolution MS/MS spectra from mass analyzers such as TOF and Orbitrap, but also to low-resolution MS/MS from ion trap devices. On the other hand, search engines that make use of high-resolution MS/MS have also been developed recently. For example, Morpheus(22) uses the specificity obtained through high-resolution mass analysis to assign charge states and remove non-monoisotopic peaks in the spectrum preprocessing stage. After such pre-processing, a very simple score is used to identify peptides, based on the sum of the number of matching product ions and the fraction of the spectrum abundance that can be assigned to matching product ions, using the target-decoy approach. Surprisingly, despite this simple approach, it was found to outperform conventional search engines. One concern with such a simplified score is that it lacks the probabilistic interpretation of the conventional scores. However, with the widespread adoption of the target-decoy approach(4), peptide-spectrum matches (PSMs) only need to be relatively ordered, and the global and local FDRs can be empirically determined. Furthermore, a Python-based open-source analysis system for large high-resolution MS datasets called AlphaPept has recently been reported(23). In this system, X! tandem score, Morpheus score or a newly introduced Generic score based on the peptide length, the total number of fragment hits, b-ion hits, and the matched intensity ratio, was used for peptide-spectrum matching, and a Percolator-like machine-learning approach was performed to maximize the peptide identification with FDR estimation. With large-scale public data now readily available, post-processor rescoring tools are becoming increasingly important(24). Furthermore, with the addition of universal spectral identifiers(25), it is becoming possible to search across projects(26). Various approaches that make use of deep learning to generate predicted MS/MS spectra and identify peptides on a large scale using the similarity of experimental data (MSBooster(27), Oktoberfest(28), MS2Rescore(29), etc.) are being developed, and an environment for integrating and re-analyzing data is gradually being established(30). However, in order to analyze these large-scale data in an integrated manner, a large-scale computing environment is often required, and there is still a need for the development of methods to facilitate the standardization and integration of large-scale data.

In this study, we propose a new, easy-to-use re-scoring method for MS/MS spectra that requires significantly less computational resources than existing tools. This scoring method, which we have named UniScore, provides a measure for selecting the most likely candidate peptide sequence from among the candidate peptide sequences, including decoy sequences, proposed by various search engines. Inspired by the PST method, we adopted the ‘number of matched amino acid sequence stretches’, in addition to the straightforward ‘total number of matched b- and y-ions’. Here, we describe the optimization of the UniScore algorithm and its application to deep global human proteome data and human phosphoproteome data registered in public repositories.

## Methods

Datasets used for UniScore development and applications were obtained from jPOST repository (https://repository.jpostdb.org/)(31) or PRIDE (https://www.ebi.ac.uk/pride/) based on ProteomeXchange identifiers, as listed in Supplementary Table 1. The LC/MS/MS raw data files and metadata used in the UniScore application were obtained from these ProteomeXchange repositories and the original publications. The multiple search engines used and their versions, the versions of the protein databases used for the search, search parameters and other information are also listed in Supplementary Table 1. For peak picking of the product ion spectra, we used the peak-picking function of MaxQuant to convert multiply charged ions to singly charged ions, declustered the isotope envelope to pick the monoisotopic peaks, recalibrated the observed *m/z* values using the pre-search results, and generated mgf files. In some cases, we used ProteoWizard(27, 32) with modifications(33) to do similar pre-processing for product ion spectra. These files were input into Comet, X! Tandem, Mascot and MaxQuant, and database searches were performed under the conditions shown in Supplementary Table 1 to obtain the respective identification results. Using the above mgf files, matching of the MS/MS spectra and peptide candidates was performed with only the top 12 peaks per *m/z* 100 Th, and the matched b- and y-ion list was output for each PSM to calculate UniScores. The entire workflow is described in Supplementary Figure 1 including a typical example of UniScore calculation. The UniScore calculation for each product ion spectrum is as follows; first, the matched b- and y-ions are counted. Based on these matched ions, the total number of amino acids flanked on both sides by these ions (called matched sequence stretch) is counted. Finally, the total number of b-ions, y-ions and matched sequence stretches are simply added together, without weighting, and the resulting value is used as the UniScore in this paper. For cases where several different sequences were assigned as candidates for a single spectrum, they were treated as chimeric spectra and the UniScore was calculated for each sequence, except when more than 50% of the annotated peaks were shared.

For the calculation of FDR, PSMs were first ordered by descending UniScore, and for each decoy hit that appeared, the decoy hit occurrence rate was calculated from the number of forward hits and decoy hits up to that score, which was then used as the FDR. When a UniScore was the same for a decoy hit and a target hit, we counted both instead of rejecting one of them. FDR at the peptide level was performed by removing redundancy in the sequence only, without considering modifications and charge, and the rest of the procedure was the same as for PSMs. For protein group FDRs, peptides were inferred to protein groups using the reported method(34), and then FDRs were obtained using the same method as above.

## Results and Discussion

As test datasets for the development of UniScore, the result files of a single-shot global proteome analysis of HeLa cells using a 2-meter-long silica monolithic column to acquire high-resolution MS/MS spectra in the DDA mode with an 8-hour gradient using Orbitrap were used. The data were acquired in five raw files, split every two hours, so that the UniScore could be evaluated independently for groups of peptides with different sets of physical properties such as precursor charge, peptide length, hydrophobicity and isoelectric point (Supplementary Figure 2). Analysis using MaxQuant enabled identification of a total of 34,746 unique peptides and 4,367 protein groups.

In connection with the development of UniScore, the parameters that can be read from the MS/MS spectrum of a matched peptide are the number of matched product ions (b, y-ion), the number of unmatched b, y-ions, the number of amino acids flanked on both sides by matched product ions (called matched sequence stretch), the peptide length of the precursor ion, the peptide mass, the fraction of the spectrum abundance that can be assigned to matched product ions, etc. The spectrum pre-processing includes top N peak abundance filtering for each bin, the determination of the charge state of the product ion, and the accumulation algorithm to the monoisotopic ions by isotope declustering.

For this test dataset, we set the initial definition of UniScore to be the sum of the number of matched product ions (b- and y-ions) and the number of matched sequence stretches, and we accepted the MaxQuant-generated peak lists with only the top 6 peaks per 50 Th bin for UniScore peak annotation. Using Mascot, MaxQuant, X! Tandem and Comet search engines, we generated UniScores and compared them with the original results from each engine. The results are shown in Figure 1A-D. In all cases for PSM hit numbers at FDR 1%, the UniScore-based approach succeeded in increasing the number of PSMs. Furthermore, by integrating the results of the four search engines using UniScore, we succeeded in maximizing the number of PSM hits at 1% FDR (Figure 1E). The correlation between the scores of each search engine and UniScore is shown in Figure 2. The coefficient of determination ranged from 0.60 (MQ andromeda score) to 0.90 (Comet Xcorr), but in all cases, a positive linear correlation was observed. We also checked the correlation between UniScore and the original score for each of the five fractions, but no change in correlation was observed with respect to physical properties such as peptide length and precursor charge (Supplementary Figure 3).

**Figure 1:**
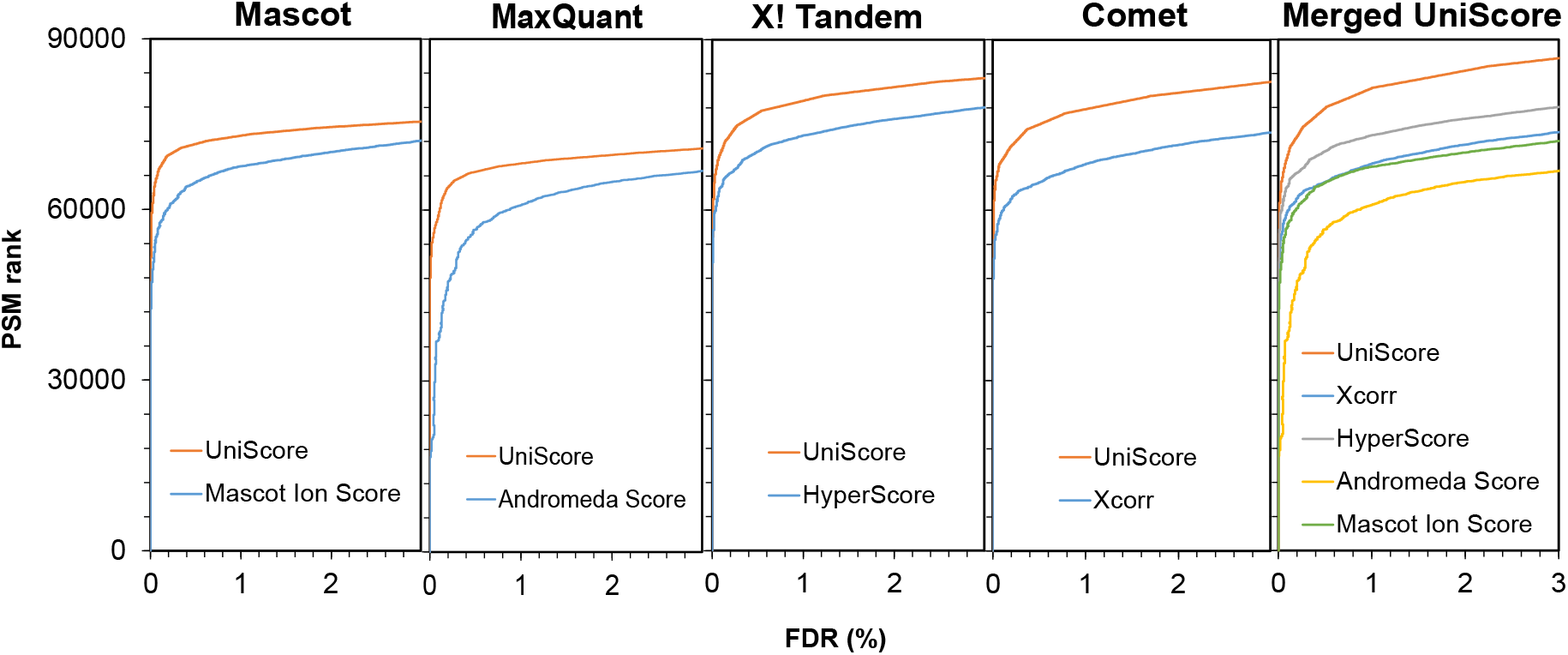
Relationship between PSM hits and FDR by the UniScore-based and the original approaches using four different search engines. The original dataset (Dataset #1) was obtained from PXD005159/JPST000200.

**Figure 2:**
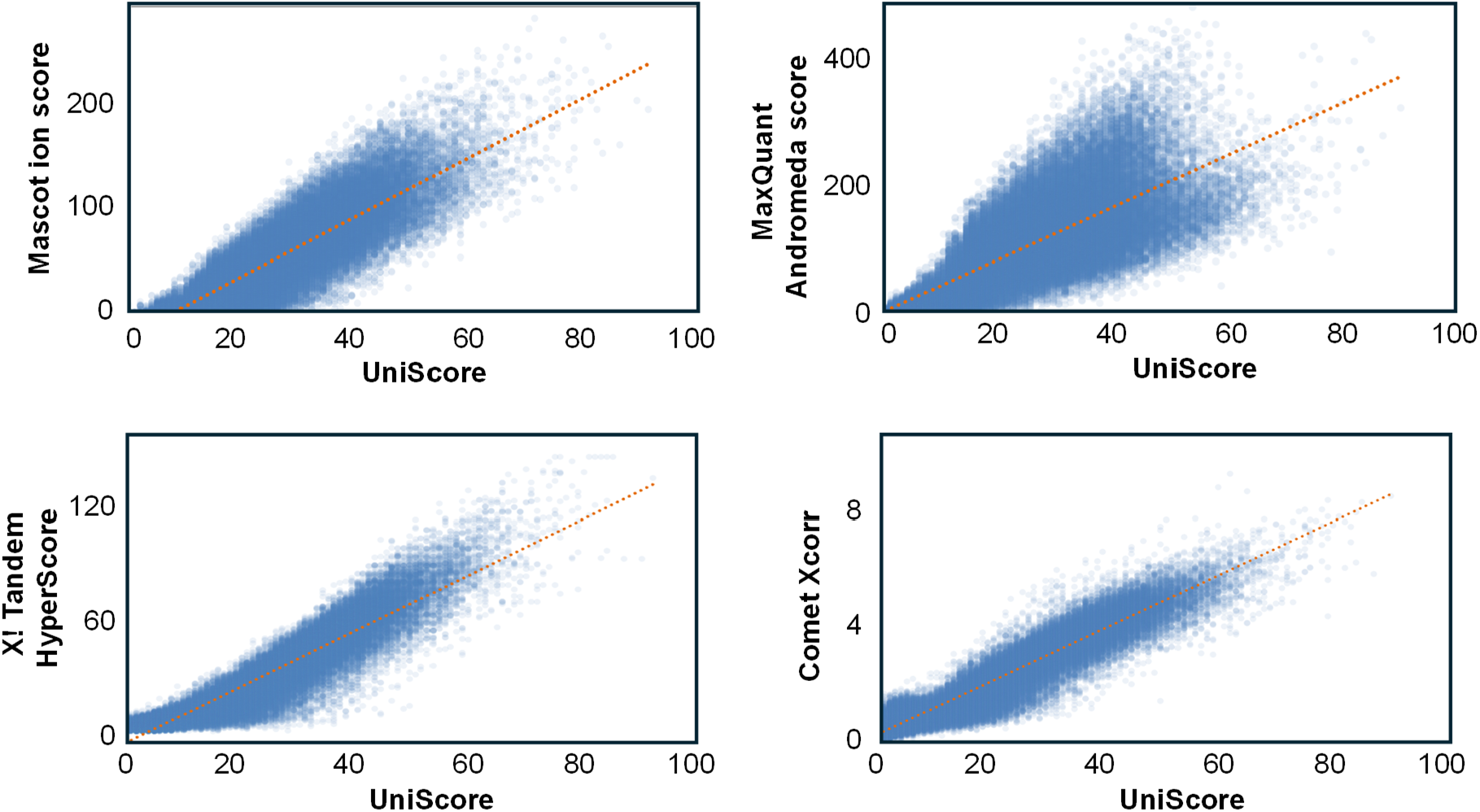
Correlation between UniScore and the scores from four different search engines including Mascot, MaxQuant, X! Tandem and Comet.

**Figure 3:**
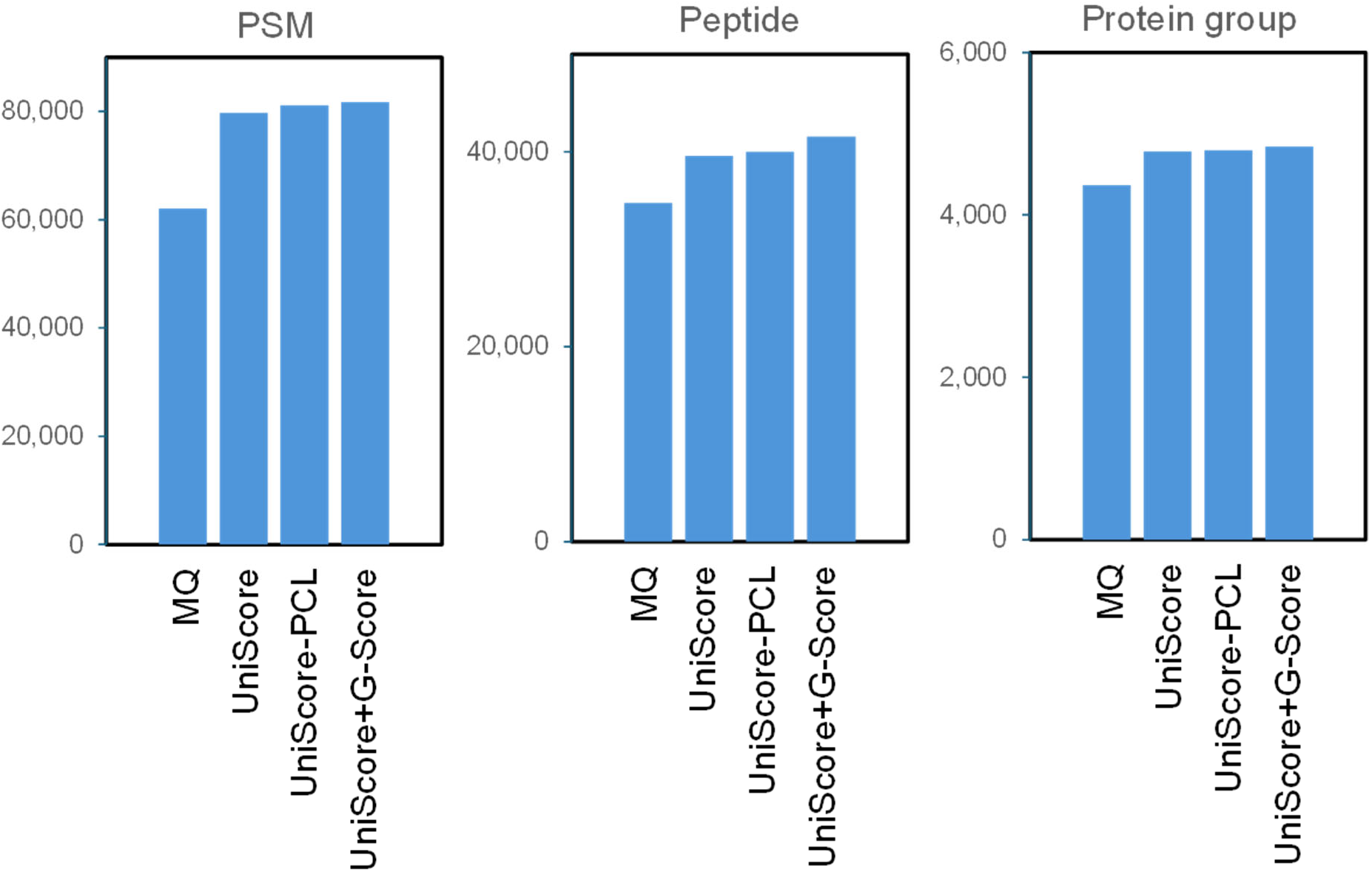
Identified numbers of PSMs, peptides and protein groups by UniScore-based approaches for Dataset #1

Next, we used ProteoWizard with the charge conversion function to create a peak list, and used Mascot to investigate the effect of converting product ions to singly-charged ions. As a result, the PSM number increased by only 2.5% with UniScore and by 3.0% with Mascot ion score, so the charge conversion effect was not significant. On the other hand, there was a clear effect of charge conversion on the UniScore value itself, and it was found that the effect of increasing the UniScore was particularly large for peptides with precursor charge numbers greater than 2 (Supplementary Figure 4). When MaxQuant was used instead of ProteoWizard as the peak list generation tool, the increase in PSM was 1.5% and 0.9% for UniScore and Mascot ion score, respectively. Since MaxQuant was slightly better as a peak list generator, it was adopted for subsequent experiments. Of the four search engines we investigated, Mascot had the lowest PSM number, and the other three are freely available, so we decided to calculate UniScore by merging the results of Comet, X! Tandem, and MaxQuant in subsequent experiments, with a view to maximizing availability.

We also considered whether to take into account the bin size, the maximum number of peaks in each bin, and the doubly charged product ions when filtering the signal intensity of product ions. As expected, we found that it was sufficient to only consider the singly charged ions, and as the bin size increased from 30, 50, to 100 Th, the optimal top N values were 4, 6, and 12, respectively. Among these, bin 100 Th, N = 12 gave the largest PSM number (Supplementary Table 2), so we decided to use this condition hereafter.

We then optimized the UniScore components and weighting. First, we considered how the number of cleavage sites identified by b- and y-ions should be counted. We considered the case where either b- or y- ions matched, and the case where both b- and y- ions matched, including weighting, and found that the coefficient of [b- or y- ion] and [b- and y- ion] was 0.5:1, that is, the maximum number of PSM was obtained when the total number of b- and y-ions was simply counted (Supplementary Table 3).

In addition, following the PST concept, the additional effect of the number of matched sequence stretches was examined, and a larger PSM was obtained when they were added together at 1:1 ratio, compared to each alone. To further examine the effect of the matched sequence stretches, we also checked the effect of the square of the number of matched sequence stretches and found a small effect (increase from 79725 to 80498 PSMs). This increase is less than 1%, so for the sake of simplicity, we decided to simply add the total number of b- and y-ions and the number of matched sequence stretches to define a UniScore. The number of unmatched b- and y-ions, the peptide length and the peptide length minus the number of matched sequence stretches were found not to improve the PSM. On the other hand, when all of the potential UniScore components such as b- or y-ions, b- and y-ions, matched sequence stretches, matched sequence stretches squared, number of unmatched b- and y-ions, peptide length minus matched sequence stretches were used as input parameters for Percolator, 81096 PSMs were obtained through machine learning-based scoring. Comparisons with the already reported Morpheus score and Generic score were also examined. The results are shown in Table 1: UniScore performed better than Morpheus, but had fewer PSMs than the Generic score. However, the use of Percolator (UniScore-PCL) slightly outperformed the number of PSMs from the Generic score. Surprisingly, the highest PSM was obtained by simply summing the UniScore and Generic score. As mentioned above, both the Morpheus score and the Generic score take into account the fraction of spectral abundance assigned to the matched product ions. Even though the contribution of this term in both scores is not very high, it is a parameter that makes the use of chimeric spectra less acceptable. It is well known that multiple peptide ions are present in the isolated window of the precursor ion in DDA measurements(35). Therefore, the UniScore + Generic score, which gives the maximum PSM, was not adopted in this study, but rather, the simple and robust UniScore or UniScore-PCL was employed for subsequent applications. Compared to the original MaxQuant results, the UniScore-based approaches increased PSMs, unique peptides and protein groups by more than 28.5 %, 14.0 % and 9.5 %, respectively.

**Table 1:**
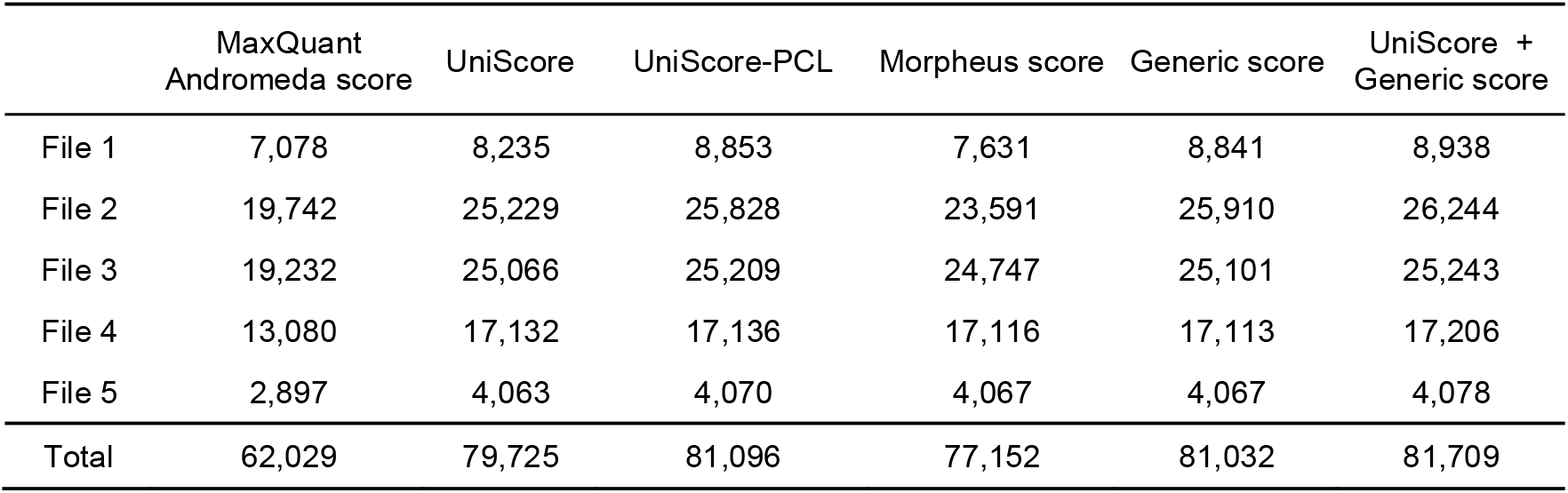
MaxQuant, UniScore and other scores for PSM at FDR 1%.

|  | MaxQuant<br>Andromeda score | UniScore | UniScore-PCL | Morpheus score | Generic score | UniScore +<br>Generic score |
| --- | --- | --- | --- | --- | --- | --- |
| File 1 | 7,078 | 8,235 | 8,853 | 7,631 | 8,841 | 8,938 |
| File 2 | 19,742 | 25,229 | 25,828 | 23,591 | 25,910 | 26,244 |
| File 3 | 19,232 | 25,066 | 25,209 | 24,747 | 25,101 | 25,243 |
| File 4 | 13,080 | 17,132 | 17,136 | 17,116 | 17,113 | 17,206 |
| File 5 | 2,897 | 4,063 | 4,070 | 4,067 | 4,067 | 4,078 |
| Total | 62,029 | 79,725 | 81,096 | 77,152 | 81,032 | 81,709 |

To confirm that the target-decoy approach appropriately controlled FDR in UniScore-based PSM identification, we performed entrapment analysis, in which the entrapment protein sequences were created from 100 randomly shuffled sequences of the *Pyrococcus furiosus* reference proteome, following the procedure used in Sage development.(36) As a result, the number of PSMs with an entrapment-based false discovery proportion (FDP) of 1% was almost the same or slightly less than the number of PSMs with an FDR of 1% using the target-decoy approach (Supplementary Table S4). Furthermore, we investigated the relationship between entrapment FDP and target-decoy FDR obtained by varying the UniScore threshold. The results showed a proportional relationship between the two, with FDP being slightly larger but almost identical (Supplementary Figure S5). These results indicate that the developed UniScore is appropriately controlled by the FDR.

Since the search engines used in this study were released quite some time ago, we compared them with the latest search engines, including Sage(36) and MSFragger(37). Furthermore, we also examined a method called “quantms,”(30) which was recently developed for the purpose of reanalyzing repository-scale data by integrating search results from different engines. The results are shown in Supplementary Table S5. These latest search engines identified more PSM than MaxQuant as expected, but not as many as UniScore. Furthermore, UniScore outperformed quantms in merging results from multiple engines, regardless of whether our original search engine set or the search engine set used in the original quantms paper(30) was employed.

As shown above, despite its simple structure, UniScore has demonstrated superior results compared to various existing statistical scores. The probability that each b/y-ion is randomly matched is reflected in the Mascot ion score(3) and Andromeda score(8), and the statistical treatment of matched sequence stretches has already been discussed in the original PST paper(20). While it may be possible to convert UniScore into a form with such statistical meaning using these factors, the current UniScore already has a proportional relationship with existing scores, as shown in Figure 2. Furthermore, if relative ranking of PSMs is possible, global and local FDRs can be empirically determined using the target-decoy approach. Therefore, it is considered impractical to convert UniScore into a statistical format. The time required for UniScore calculation is shown in Supplementary Table S6. UniScore calculations are extremely fast, even with large data files. This means that large-scale re-scoring can be performed without the need for massive computing resources.

### Peptide-Chimeric Spectrum Matching

There is a possibility that the peptide sequences suggested by the three search engines will differ for a single spectrum, and this is due to the simultaneous isolation of different precursor ions, where product ions of different origins coexist in a single MS/MS spectrum (chimeric spectrum). The precursor ion fraction in standard DDA is reported to be 0.14 (35), so it is thought that chimeric spectra are obtained with a very high probability. The total number of annotated spectra in this test data set was 68,829, and the number of chimeric spectra was 8,676. The number of PSMs per spectrum was 2 for most of the spectra, and the number of spectra with 3 or more PSMs was only 0.7% (Figure 4A). We then investigated the distribution of the overlap rate of annotated product ions in chimeric spectra with product ions from other precursors (Figure 4B). More than half of the spectra had no overlap at all, and there were almost no cases where the overlap rate exceeded 50%. This suggests that when the search engines returned the sequence candidates, those with a high overlap rate for the product ions had already been removed, and only the best hits per spectrum remained as candidate sequences. These results indicate that the number of chimera spectrum-derived PSMs in the case of UniScore is acceptable without additional criteria.

**Figure 4:**
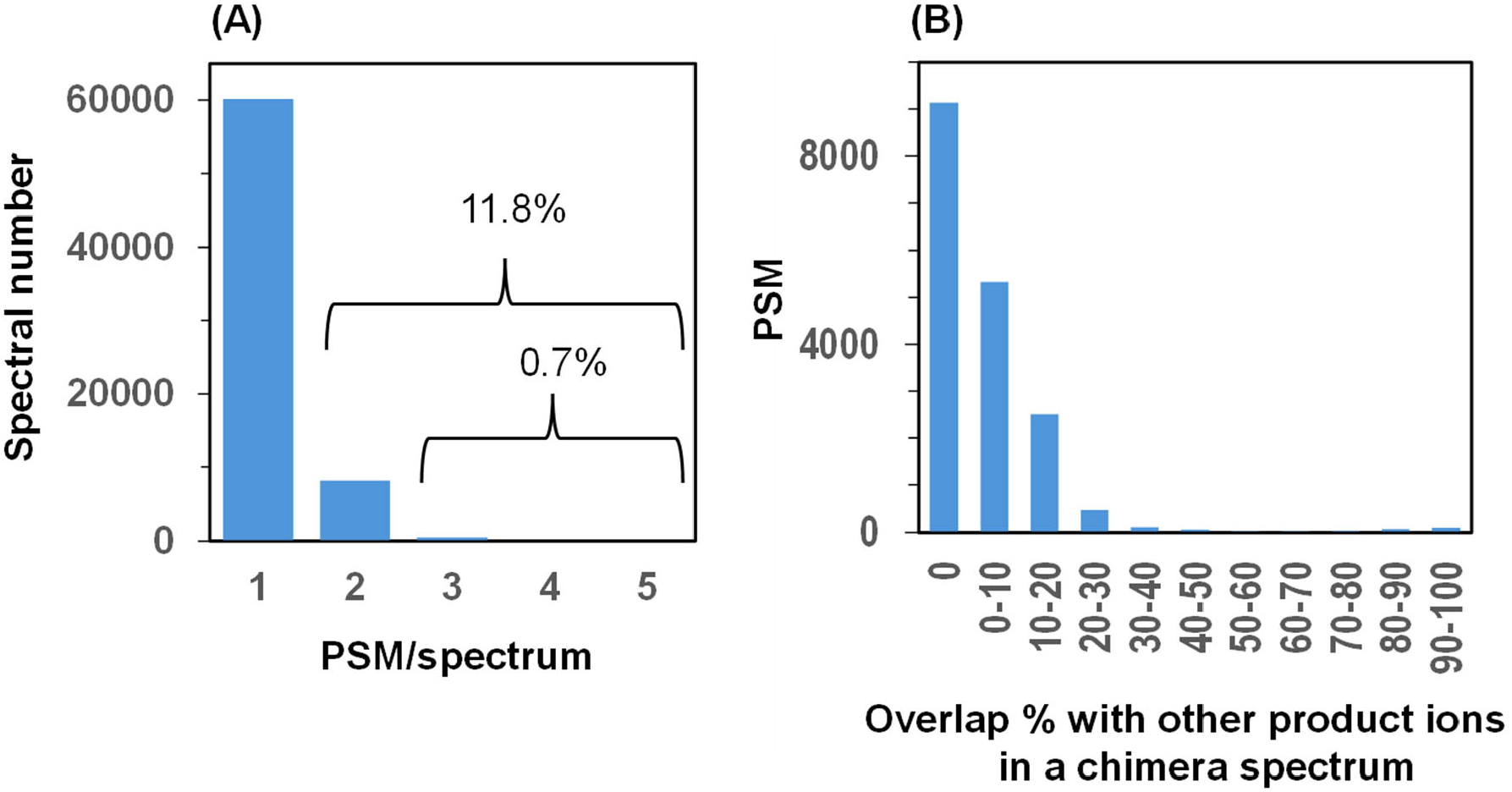
PSMs in chimeric spectra. (A) Distribution of PSM number per spectrum, (B) Overlap percentage of product ions with those from other precursor ions in a chimeric spectrum.

### Application to human proteome data by LC/MS/MS with extensive pre-fractionation

We selected the HeLa proteome analysis data by Bekker-Jensen et al.(38) as an example of how a combination of high-pH RP fractionation and a short gradient can achieve both high efficiency and high depth, and we re-analyzed their data using UniScore. In the original paper, they analyzed 1 µg of HeLa peptides on the column from each of 46 HpH fractions with 30 min LC/MS/MS gradients (45 min run-to-run), taking a total of 34.5 hr, and this resulted in identification of 166,620 unique peptide sequences covering 11,292 protein groups or 10,284 protein-coding genes according to MaxQuant. In this study, we re-analyzed the data using MaxQuant, UniScore and UniScore-PCL with the same database and search conditions. Fig. 5A shows a comparison of the PSM values for each of the 46 fractions. In all fractions, the UniScore approach outperformed MaxQuant by about 20% in terms of PSM values, and UniScore-PCL gave slightly better results than UniScore. After merging all the results from the 46 fractions, we calculated the number of identified PSMs, peptides, and protein groups at FDR 1% by UniScore and compared them with the MaxQuant results (Figure 5B,C,D). As expected, UniScore outperformed MaxQuant, but in this case the difference was not as great as in the results from the Dataset #1 used in algorithm development. This is because it has already been shown that increasing the number of human protein identifications by 1000 from 9000 is completely different from increasing it by 1000 from 4000.(39) The UniScore-based chimera spectrum content for this dataset was 5.6%, less than half the value of 11.8% for the Dataset #1. This is because the Dataset #1 was obtained in a single shot, whereas this dataset was heavily fractionated.

**Figure 5:**
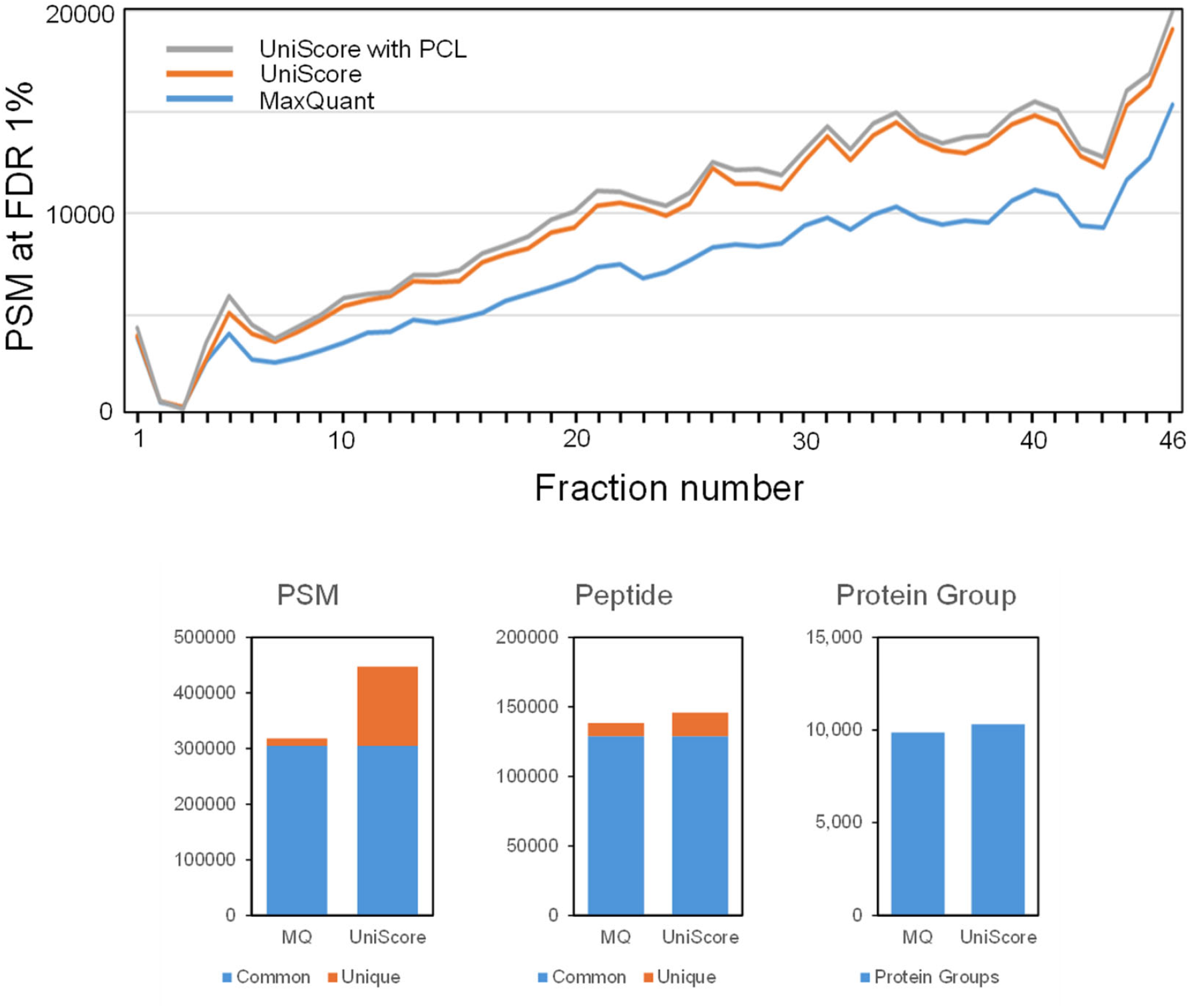
Deep HeLa proteome data analyzed by MaxQuant and UniScore.

### Application to human phosphoproteome data obtained by LC/MS/MS following phosphopeptide enrichment

As an example of application to post-translational modification proteome data, we used UniScore to examine phosphorylation proteome data(40) obtained from Caco-2 cells infected with SARS-CoV-2. In that work, TMT labeling was performed, followed by phosphopeptide enrichment using Fe-IMAC, and high-pH RP fractionation was performed using StageTip, followed by LC/MS/MS analysis. Proteome Discoverer implemented with Sequest HT was used for data analysis, and the data were controlled at 1% FDR at the peptide level using Percolator with the TD method. Phosphorylation site localization was determined using PhosphoRS implemented within Proteome Discoverer. As we did not have this commercial tool for determining phosphosite localization, we applied 1% FDR at the peptide level, removed TMT unlabeled peptides, and then compared only the sequences and number of phosphate groups without considering the localization of phosphosites. The results are shown in Figure 6. The UniScore approach significantly improved the number of identified phosphopeptides by 73.2%. When we investigated what kind of phosphorylated peptides were specifically identified by UniScore, we found that longer phosphopeptides were being identified. One concern with UniScore is that, due to its definition formula, it tends to give higher scores to longer peptides. However, as already shown in Supplementary Figure 3, the bias towards long peptides is almost the same as in the Mascot ion score, which is scored based on the probability of random matching. In other words, both UniScore and Mascot ion score tend to be high for long peptides. Nevertheless, it should be noted that this is not a statistical bias, because the database search space for precursor ions with higher molecular mass is also narrowed. In any case, UniScore was shown to be effective, particularly for phosphopeptides, which tend to have a long average peptide length.

**Figure 6:**
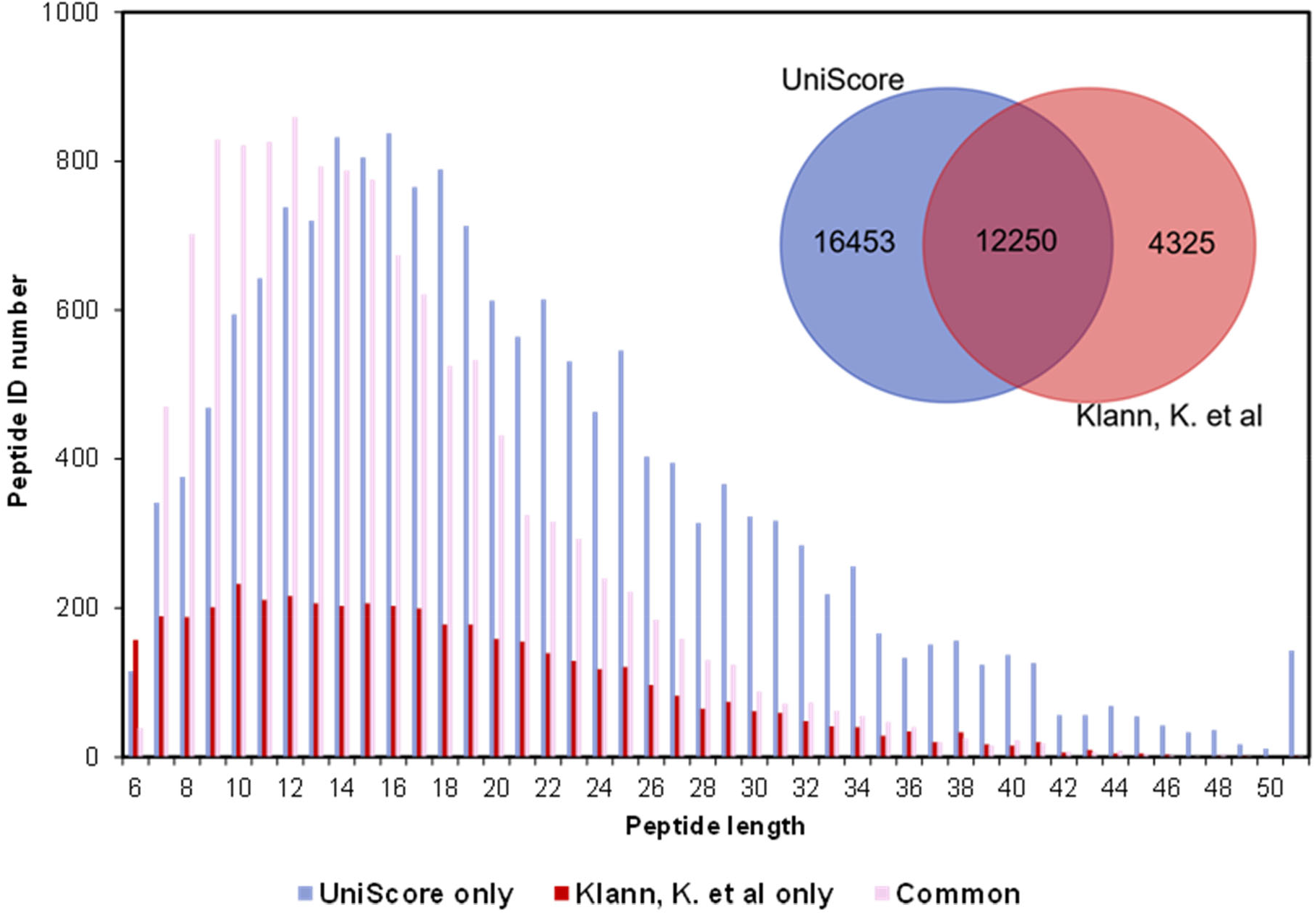
UniScore re-analysis of phosphoproteome data from SARS-CoV-2-infected Caco-2 cells. The Venn diagram shows the identification results from the original paper and by the UniScore approach for identified phosphopeptides. The bar graph shows the length distribution of peptides uniquely identified by each method and commonly identified peptides. Note that phosphosite localization was not considered for this analysis.

## Conclusions

UniScore can be computed by using only the product ion information matching the sequence, making it an easy tool that requires significantly less computational resources than various existing rescoring tools. The combination of multiple search engines to output candidate sequences without bias has not yet been fully optimized. Nevertheless, the large gain over each of the examined search engines indicates the potential value of this approach. In matching against chimera spectra, the number of PSMs per spectrum is also well-balanced and not too large. In principle, matching with the predicted MS/MS spectra using deep learning should be more powerful for maximizing the identification number, but since the MS/MS spectrum itself is dependent on experimental conditions, the prediction accuracy is dependent on factors such as the collision energy and the characteristics of the model. UniScore, on the other hand, is a robust measure that is virtually unaffected by experimental conditions and should be more widely applicable. Further studies are needed to confirm its robustness (repository scale analysis) by applying it to larger and more diverse data sets (various post-translational modifications, coexistence of Orbitrap and Q-TOF data, etc.). This UniScore concept can also be extended to data-independent acquisition data in which prioritized selection of product ions would be one of the key steps. Further development is on-going in our laboratory.

## Supporting information

all supple figures and tables

## DATA AVAILABILITY

The UniScore re-analysis results for the Datasets #1, #2 and #3 are available through the jPOST repository (https://jpostdb.org/) as RPXD034294, RPXD056882, and RPXD034696, respectively. The UniScore program is also available in GitHub (https://github.com/jPOST-tools/UniScorePrograms20241016) with the data input template.

### ACKNOWLEDGEMENTS

We thank all members of the jPOST project team. This work was supported by the Life Science Database Integration Project (Database Integration Coordination Program) funded by the Office of the NBDC Program of the Japan Science and Technology Agency [18063028, JPMJND2304].

## CONTRIBUTIONS

Y.I. developed the initial concept. T.T. executed computational experiments, T.T., A.C.Y., K.O., C-H.C., N.A., N.S. and Y.I. carried out data analyses. Y.I. supervised the project and provided resources for the study. Writing—review and editing was carried out by Y.I. Approval of the final version of the paper was carried out by all authors.

## ETHICS DECLARATIONS

All authors declare no competing interests.

