## Supplementary material for "UniScore, a unified and universal measure for peptide identification by multiple search engines": all supple figures and tables

---

### Table of contents

**Figure S1.** UniScore calculation

**Figure S2.** Base-peak chromatogram of a HeLa global proteome sample.

**Figure S3.** Correlation between UniScore and Mascot ion score for peptides with different physical properties in Files 1-5.

**Figure S4.** Effect of charge conversion on UniScore of peptides with different precursor charges. The red dotted line shows y = x.

**Figure S5.** Decoy-based FDR using linear discriminant analysis is well-calibrated against entrapment-based FDP.

**Table S1.** Datasets and data analysis parameters used in this study.

**Table S2.** Effect of product ion abundance filtering on UniScore-PSM at FDR 1%.

**Table S3.** Effect of matched b-, y-ions on UniScore-PSM at FDR 1%.

**Table S4.** UniScore-based PSMs at FDR 1% (target-decoy) and FDP 1% (entrapment).

**Table S5.** Various Search Engines, UniScore and Quantms for PSM at FDR 1%.

**Table S6.** Estimation of UniScore Calculation

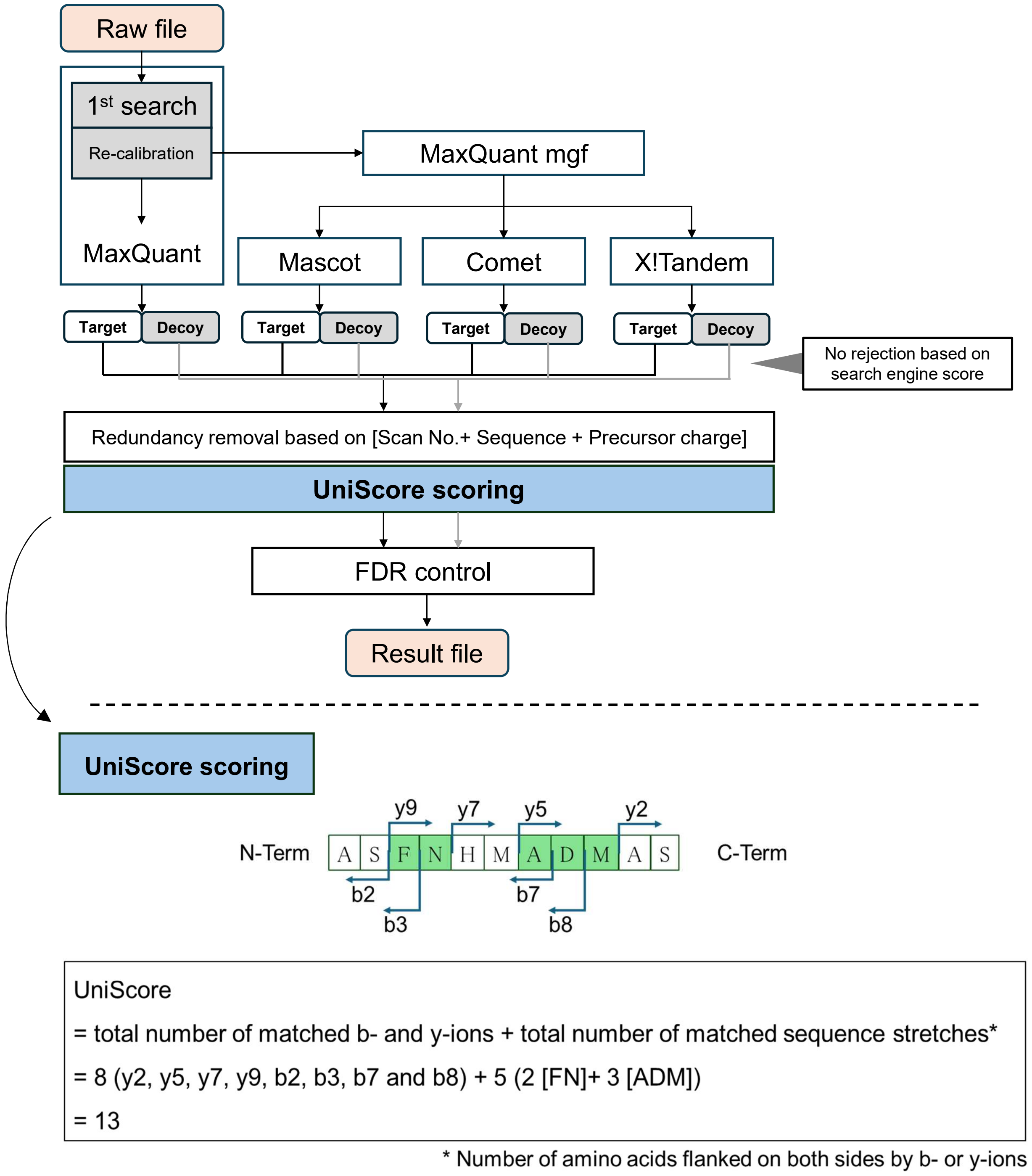


**Supplementary Figure S1:** UniScore calculation

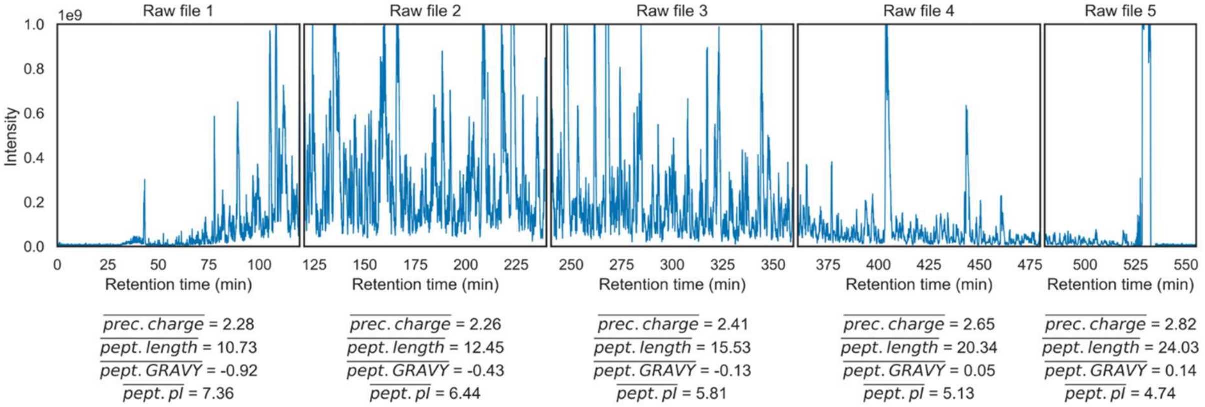


**Supplementary Figure S2**: Base-peak chromatogram of a HeLa global proteome sample. Data was obtained from PXD005159/JPST000200.

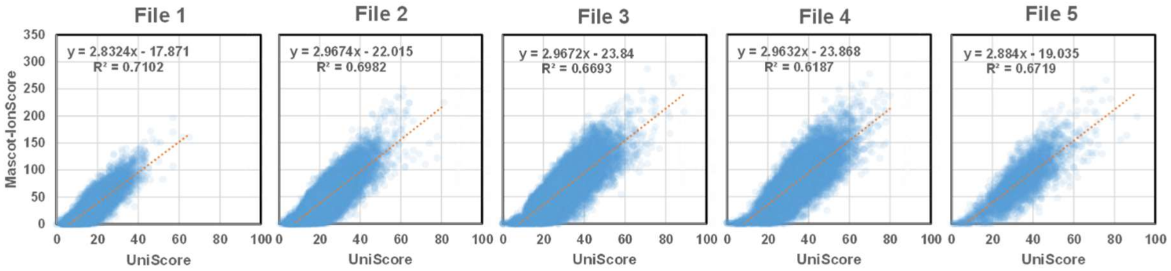


**Supplementary Figure S3**: Correlation between UniScore and Mascot ion score for peptides with different physical properties in Files 1-5.

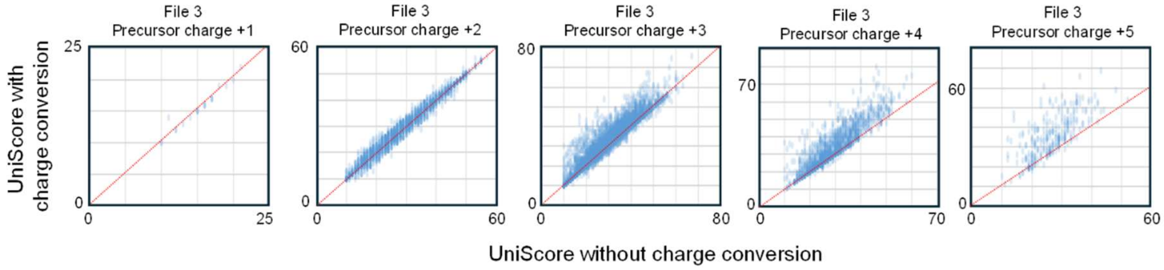


**Supplementary Figure S4**: Effect of charge conversion on UniScore of peptides with different precursor charges. The red dotted line shows y = x.

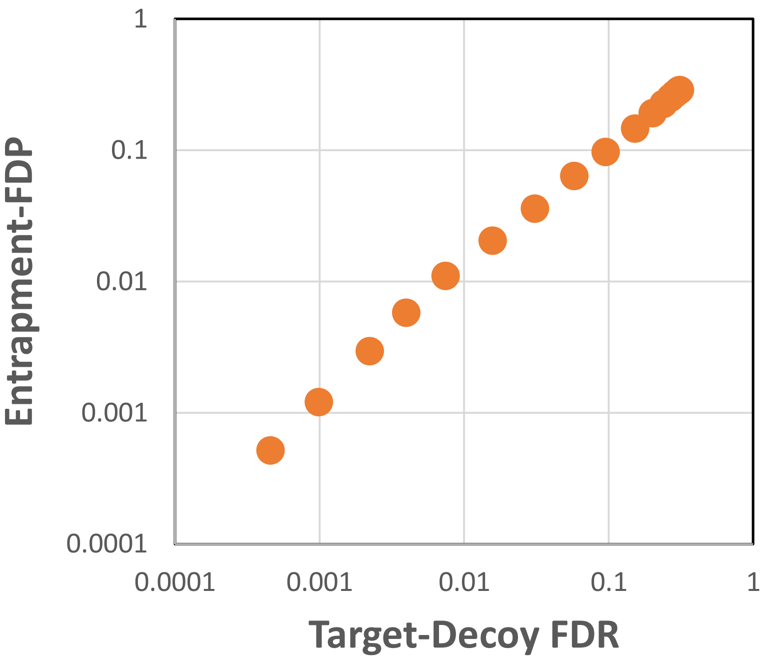


**Supplementary Figure S5**: Decoy-based FDR using linear discriminant analysis is well-calibrated against entrapment-based FDP.

**Supplementary Table S1:** Datasets and data analysis parameters used in this study

| **Datasets** | | |
|---|---|---|
| Dataset #1<br>for UniScore development | Original ID | PXD005159 |
| | Reanalysis ID | RPXD034294/ JPST001624 |
| Dataset #2<br>for UniScore application<br>(global proteomics) | Original ID | PXD004452 |
| | Reanalysis ID | RPXD056882/ JPST003443 |
| Dataset #3<br>for UniScore application<br>(phosphoproteomics) | Original ID | PXD018357 |
| | Reanalysis ID | RPXD034696/ JPST001780 |
| **Search engines and other tools with version** | | |
| Mascot | 2.7.0 | |
| MaxQuant: | 1.6.17.0（Download date: 2021/01/26） | |
| Comet | 2019.01 rev. 5 | |
| X! Tandem | 2015.04.01.1 | |
| Percolator | v3-04 | |
| ProteoWizard | 3.0.21021.e2078b75f 64-bit（Download date:2021/01/25） | |
| **Reanalysis search parameters** | | |
| Database<br>Enzyme<br>Fixed modifications<br>Variable modifications<br>MS1 tolerance<br>MS2 tolerance<br>Missed cleavage | Listed in jPOST repository (https://repository.jpostdb.org/) with RPXD identifiers | |
| Input peak list | mgf files generated by MaxQuant | |
| Peak matching for UniScore | Top 12 peaks within every 100 Th bin | |

### **Supplementary Table S2:** Effect of product ion abundance filtering on UniScore-PSM at FDR 1%

| Bin size (Th) | 30 | | | 50 | | | | | | | 100 | | | | | | | | |
|---|---|---|---|---|---|---|---|---|---|---|---|---|---|---|---|---|---|---|---|
| Top N/bin | 3 | 4 | 5 | 4 | 5 | 6 | 6 | 7 | 8 | 10 | 6 | 8 | 10 | 11 | **12** | 12 | 13 | 14 | 15 |
| Product ion charge | 1 | 1 | 1 | 1 | 1 | 1 | 1,2 | 1 | 1 | 1 | 1 | 1 | 1 | 1 | **1** | 1,2 | 1 | 1 | 1 |
| File 1 | 7,901 | 8,253 | 7,972 | 8,199 | 8,084 | 8,266 | 7,909 | 7,911 | 7,995 | 8,080 | 7,654 | 7,765 | 8,114 | 8,228 | **8,296** | 7,961 | 8,347 | 8,383 | 8,404 |
| File 2 | 23,257 | 23,769 | 23,445 | 22,995 | 23,530 | 23,767 | 23,823 | 23,898 | 23,465 | 23,576 | 22,048 | 23,189 | 23,669 | 23,768 | **23,840** | 23,353 | 23,350 | 23,411 | 23,446 |
| File 3 | 21,898 | 21,862 | 21,887 | 21,853 | 21,926 | 21,943 | 21,874 | 21,879 | 21,889 | 21,897 | 21,739 | 21,882 | 21,926 | 21,935 | **21,947** | 21,875 | 21,873 | 21,879 | 21,885 |
| File 4 | 14,400 | 14,404 | 14,408 | 14,390 | 14,399 | 14,405 | 14,410 | 14,407 | 14,408 | 14,409 | 14,383 | 14,397 | 14,401 | 14,405 | **14,405** | 14,409 | 14,406 | 14,406 | 14,408 |
| File 5 | 3,290 | 3,290 | 3,290 | 3,290 | 3,290 | 3,291 | 3,292 | 3,291 | 3,291 | 3,291 | 3,288 | 3,288 | 3,289 | 3,290 | **3,291** | 3,292 | 3,291 | 3,291 | 3,291 |
| Total | 70,746 | 71,578 | 71,002 | 70,727 | 71,229 | 71,672 | 71,308 | 71,386 | 71,048 | 71,253 | 69,112 | 70,521 | 71,399 | 71,626 | **71,779** | 70,890 | 71,267 | 71,370 | 71,434 |

Red bold numbers indicate the maximum number of PSMs in each row. Search parameters as listed in Supple Table 1 except the parameters in top N in bin X Th

### **Supplementary Table S3:** Effect of matched b-, y-ions on UniScore-PSM at FDR 1%

| Coefficient for (b and y) sites x 2 | 0 | 0.1 | 0.2 | 0.3 | 0.4 | **0.5** | 0.6 | 0.7 | 0.8 | 0.9 | 1 |
|---|---|---|---|---|---|---|---|---|---|---|---|
| Coefficient for (b or y) sites | 1 | 0.9 | 0.8 | 0.7 | 0.6 | **0.5** | 0.4 | 0.3 | 0.2 | 0.1 | 0 |
| File 1 | 1,270 | 1,616 | 3,078 | 5,241 | 7,442 | **7,544** | 6,067 | 4,439 | 3,650 | 3,528 | 3,143 |
| File 2 | 8,359 | 11,315 | 15,403 | 19,413 | 23,195 | **23,426** | 20,988 | 18,296 | 15,158 | 14,603 | 10,723 |
| File 3 | 14,576 | 17,022 | 20,985 | 22,758 | 24,254 | **24,687** | 23,752 | 21,676 | 18,229 | 16,598 | 14,582 |
| File 4 | 14,629 | 16,280 | 16,848 | 17,137 | **17,161** | 17,143 | 16,958 | 16,301 | 14,927 | 13,253 | 11,023 |
| File 5 | 3,931 | 4,021 | 4,053 | **4,073** | 4,069 | 4,061 | 4,020 | 3,824 | 3,200 | 2,333 | 1,890 |
| Total | 42,765 | 50,254 | 60,367 | 68,622 | 76,121 | **76,861** | 71,785 | 64,536 | 55,164 | 50,315 | 41,361 |

Red bold numbers indicate the maximum number of PSMs in each row.

**Supplementary Table S4**: UniScore-based PSMs at FDR 1% (target-decoy) and FDP 1% (entrapment).

| | UniScore | |
|---|---|---|
| | Target-decoy FDR 1% | Entrapment FDP 1% |
| File 1 | 8,235 | 8,853 |
| File 2 | 25,229 | 25,828 |
| File 3 | 25,066 | 25,209 |
| File 4 | 17,132 | 17,136 |
| File 5 | 4,063 | 4,070 |
| Total | 79,725 | 81,096 |

**Supplementary Table S5**: Various Search Engines, UniScore and Quantms for PSM at FDR 1%.

| | MaxQuant | MSFragger 4.1 | Sage 0.14.7 | MaxQuant/Comet/XTandem/Mascot | | | Comet/Sage/MSGF+ | | |
|---|---|---|---|---|---|---|---|---|---|
| | | | | UniScore | UniScore-PCL | Quantms | UniScore | UniScore-PCL | Quantms |
| File 1 | 7,078 | 8,253 | 8,070 | 8,235 | 8,853 | 7,097 | 6,988 | 8,192 | 6,857 |
| File 2 | 19,742 | 21,274 | 21,593 | 25,229 | 25,828 | 22,031 | 23,545 | 25,183 | 20,070 |
| File 3 | 19,232 | 20,209 | 20,290 | 25,066 | 25,209 | 22,301 | 24,395 | 24,933 | 21,968 |
| File 4 | 13,080 | 14,339 | 14,368 | 17,132 | 17,136 | 15,651 | 16,589 | 16,747 | 15,016 |
| File 5 | 2,897 | 3,424 | 3,503 | 4,063 | 4,070 | 3,837 | 3,865 | 3,900 | 3,185 |
| Total | 62,029 | 67,499 | 67,824 | 79,725 | 81,096 | 70,917 | 75,382 | 78,955 | 67,096 |

**Supplementary Table S6**: Estimation of UniScore Calculation

- CPU — Intel Xeon Gold 6248R @ 3.00GHz
- memory — 512GB
- OS — Windows Server 2016 Standard
- programming language — Ruby 2.7 x64
- UniScore calculation process — Single thread coding

| ID | File name | Raw file size | mgf size | UniScore computation time (h:m:s) |
|---|---|---|---|---|
| JPST000200 | 150211tk04-whole_2m8h-1.raw | 1.3G | 160M | 0:01:50 |
| | 150211tk04-whole_2m8h-2.raw | 1.1G | 269M | 0:04:14 |
| | 150211tk04-whole_2m8h-3.raw | 974M | 194M | 0:03:12 |
| | 150211tk04-whole_2m8h-4.raw | 681M | 111M | 0:02:02 |
| | 150211tk04-whole_2m8h-5.raw | 578M | 40M | 0:00:31 |
